# Satb1integrates DNA sequence,shape, motif densityand torsional stressto differentially bind targets in nucleosome-dense regions

**DOI:** 10.1101/450262

**Authors:** Rajarshi P. Ghosh, Quanming Shi, Linfeng Yang, Michael P. Reddick, Tatiana Nikitina, Victor B. Zhurkin, Polly Fordyce, Timothy J. Stasevich, Howard Y. Chang, William J. Greenleaf, Jan T. Liphardt

## Abstract

Satb1 is a genome organizer that regulates multiple cellular and developmental processes. It is not yet clear how Satb1 selects different sets of targets throughout the genome. We used live-cell single molecule imaging and deep sequencing to assess determinants of Satb1 binding-site selectivity. We found that Satb1 preferentially targets nucleosome-dense regions and can directly bind consensus motifs within nucleosomes. Some genomic regions harbor multiple regularly spaced Satb1 binding motifs (typical separation ∼1 turn of the DNA helix), characterized by highly cooperative binding. The Satb1 homeodomain is dispensable for high-affinity binding but is essential for specificity. Finally, Satb1⇔DNA interactions are mechanosensitive: increasing negative torsional stress in DNA enhances Satb1 binding and Satb1 stabilizes base unpairing regions (BURs) against melting by molecular machines. The ability of Satb1 to control diverse biological programs may reflect its ability to combinatorially use multiple site selection criteria.

## Introduction

At the core of cellular information processing is the ability of transcription factors (TFs) to bind subsets of genomic targets selectively. Proteins with multiple DNA Binding Domains (DBD) can combinatorially engage multiple distinct core DNA consensus motifs.^1,2^ The DNA backbone and major and minor groove shape constitute a second evolutionarily conserved constraint^3^ recognized through “indirect readout”^4^ or “shape readout” ^5,6^ mechanisms. Recently it has been suggested that dinucleotide features are sufficient to reliably predict DNA shape parameters ^7^. Transcription factors have also been shown to streamline their binding choices based on the shape of regions flanking the core motif.^8^ Beyond sequence and shape, geometric constraints such as motif-to-motif spacing, orientation, and local density of binding sites influence cooperative TF binding.^9,10^ Finally, TF binding is further modulated by chromatin accessibility and nucleosome occupancy^11,12^ which in turn are affected by torsional stress and DNA deformability.^13–14,15,16,17^ Our goal was to explore how the interplay of these parameters defines the genome-wide distribution of the chromatin “loopscape” regulator^18^ Special AT-Rich Sequence Binding Protein 1 (Satb1).

Satb1 is a dimeric/tetrameric^19^ transcription factor with multiple DNA binding domains, namely CUT1, CUT2 and a C-terminal homeodomain (HD). Satb1 has been implicated in diverse cellular processes including epidermal differentiation^20^, breast cancer metastasis^21^, thymocyte development^22^, Th2 cell activation and cytokine production^23^, cortical development^24^, X-chromosome inactivation^25^, and embryonic stem cell differentiation.^26^ Satb1 has also been deemed to be a “genome organizer” responsible for rapid phenotypic transitions.^18^ For instance, 3C techniques applied to T cells suggest that Satb1 mediates transcriptional and epigenetic changes at target loci.^23^ In the same vein, Satb1 is instrumental in establishing a Foxp3^+^ regulatory T cell (T_reg_)-specific lineage by defining a T_reg_ cell-specific super enhancer landscape.^27^

Despite an abundance of *in vitro* biochemical and biophysical data, there is no consensus on Satb1’s binding mechanism(s). *In vitro* and affinity-based pull-down experiments indicate that Satb1 binds A/T rich motifs with high base-unpairing potential (BURs)^28,29^, which have a propensity to unpair under torsional stress.^30,31^ Chemical interference assays suggest that Satb1 binds along the minor groove of DNA with virtually no contact with the bases.^32,28^ SELEX experiments suggest that the Satb1 homodimer binds an inverted AT-rich palindromic repeat along the minor groove.^32^ Unlike other HD proteins, Satb1’s HD does not independently bind an HD consensus sequence but appears to increase binding specificity towards BURs *in vitro.*^29^ It has also been postulated that the spacing of the half-sites is critical for binding as a dimer. Finally, solution biophysical assays suggest that the sequence specificity is due to binding by CUT1 along the major groove^33,34^ but does not require an AT-rich inverted palindromic repeat.^34^ The disparate nature of these findings could reflect over-simplified *in vitro* biochemical settings (which exclude the native genomic environment and/or nucleosomal context) or the use of a narrow subset of DNA substrates with selection bias.

To elucidate how Satb1 binds its targets, we combined live cell imaging (spatiotemporal FRAP^35^, image correlation spectroscopy^36^, and single molecule tracking^37,38^), genomics (ChIP-seq^39^, ATAC-seq^40^, ChIP-ATAC-seq, TMP-seq^15^) and *in vitro* nucleosome binding assays. We found that Satb1 binds transposase-inaccessible, nucleosome-dense regions in chromatin and contacts motifs embedded in nucleosomal core sequences, both *in vitro* and *in vivo*. Satb1 binding showed repeated binding and unbinding to a small number of spatially proximal chromatin interaction hotspots. Deep sequencing revealed clusters of Satb1 binding sites in the genome, characterized by highly cooperative binding. Satb1 preferred regions of the DNA with multiple consensus motifs spaced ∼10 bp apart. The HD was crucial for site-specificity, as it helped to home in on motifs with flanks having enhanced negative propeller twist and higher AT content than the genome average. Finally, Satb1 showed more efficient binding to target sites under highly negative torsional stress and stabilized base-unpairing regions (BURs) against helix destabilization, as evidenced by the abolition of psoralen crosslinking from binding sites in the presence of Satb1.

## Results and Discussion

### Satb1 distribution and dynamics in live cells reveals chromatin interaction hotspots

In thymocytes, the pool of nuclear Satb1 appears to be concentrated in extended ‘tendrils’ which may be important for thymocyte development.^18^ To investigate whether the Satb1 molecules in these extended regions are organized into a static structure of some kind or if these tendrils represent an excluded volume of dynamically exchanging Satb1, we tagged native Satb1 with eGFP using a CRISPR/CAS9 knockin strategy in an immortalized CD4+CD8+thymocyte cell line (VL3-3M2, Supplementary Figure 1a, b). Transient co-expression of histone H2B-mCherry revealed an inverse spatial correlation of Satb1 and dense heterochromatin, confirming previous immunofluorescence studies^23^ (Figure 1a, b), even though the levels of Satb1 in heterochromatin were clearly detectable (Figure 1a, b).

**Figure 1.**
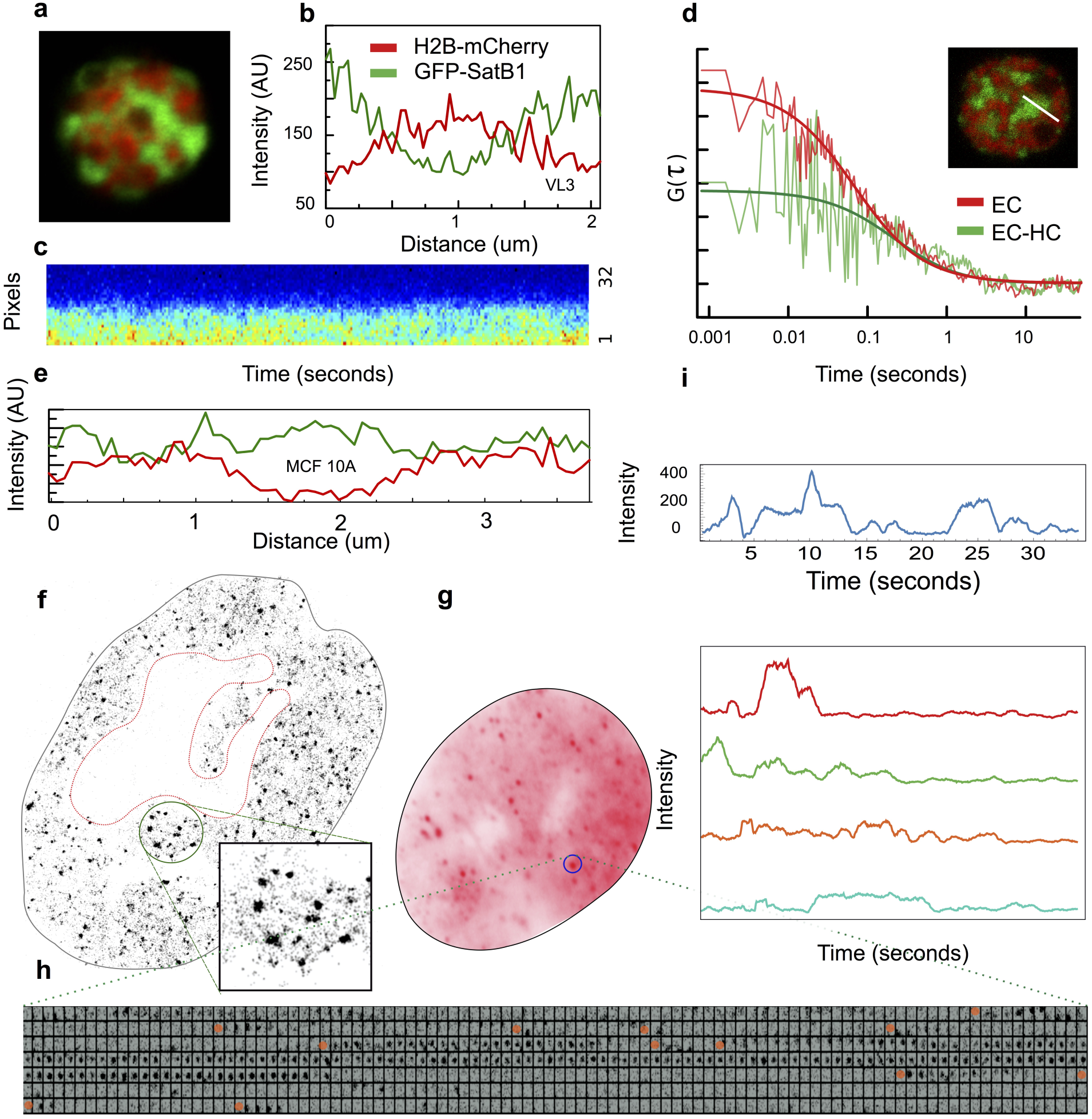
**High resolution spatiotemporal analysis of SatB1 binding in live cells. a**. A VL3 3M2 thymocyte Satb1-eGFP knockin cell line co-expressing core histone H2B-mCherry shows the classic “cage pattern”. **b**. Satb1-eGFP and H2B-mCherry profile along a line bisecting the thymocyte nucleus shows smaller amount of Satb1 in regions with high H2B-mCherry signal. **c, d**. A 3.2 μm-line bisecting a Satb1 cage and an adjacent heterochromatin is scanned repeatedly with a pixel dwell time of 6.3 μs (32 pixels) and a line time of 0.47 ms. **c**. A fluorescence intensity carpet generated by repeated scanning over time where the pixels are along y axis and time is along x axis. **d**. The autocorrelation function is calculated pixel-by-pixel along the scanned line. The two autocorrelation functions shown are for one-pixel columns that reside either in the cage or at the cage-heterochromatin boundary. **e**. Exogenous Satb1 distribution in MCF-10A cells does not show any stark difference between euchromatic and heterochromatic compartments. **f.** Super-resolution image of Satb1 in the MCF-10A nucleus reconstructed from all localizations obtained over ∼1000 frames. **g**. Properties of the Satb1 interaction hotspots. The summed intensity of 700 frames (spanning 35 seconds) for a cell expressing eGFP fusion of native Satb1. Note discrete spot like structures scattered throughout the nucleus. **h**. A 7x100 matrix of one region of interest (marked by circle in **f**) showing intensity changes over time. **i**. Intensity traces over time of a few chromatin interaction hot spots.

We used image correlation spectroscopy (ICS) to measure the dynamics of Satb1-eGFP in the euchromatin as well as at the euchromatin/heterochromatin (EC/HC) boundary. Figure 1c shows a carpet of pixel intensity along a 3.2 μm line bisecting a Satb1 ‘tendril’ vs. time. Temporal autocorrelation analysis^36^ of intensities of each pixel along the bisecting line yielded diffusion coefficients of Satb1 specific to that pixel (Materials and Methods, Figure 1d). The diffusion coefficients obtained from the average of all fits showed that Satb1 is highly dynamic in EC (5.1±1.8 μm^2^/s) and to a lesser extent at the EC/HC boundary (1.7±1.1 μm^2^/s). Therefore, the majority of the Satb1 molecules are highly dynamic albeit with varying ability to access different regions of the nucleus, raising the possibility that Satb1 ‘tendrils’ are not static structures in their own right, but reflect cell-type specific features of the thymocyte heterochromatin.

We then used HiLo TIRF microscopy^41^ to localize single, chromatin-bound Satb1 molecules. To set a baseline and assess how Satb1 binds to virgin chromatin, we ectopically expressed a C terminal eGFP fusion of Satb1 (inducible via cumate control; Material and Methods) in the MCF10A breast epithelial cell line. MCF10A lacks native Satb1. In this experimental geometry, the Satb1-eGFP will bind a genome that is unaffected by architectural changes and epigenetic modifications resulting from chronic Satb1 exposure. Unlike thymocytes which exhibit a specialized nuclear architecture, the Satb1 distribution in the MCF10A did not show marked depletion in heterochromatin (Figure 1e, Supplementary Figure 1c). The background (leaky) zero-cumate expression yielded a molecule observation density of ∼30 per nucleus/frame in HILO TIRF imaging^41^, which proved suitable for single molecule tracking.

Localization of single Satb1 molecules (Materials and Methods, Figure 1f) revealed a wide spatial variation in signal density, ranging from regions with only a few localizations per μm^2^ to ‘hotspots’ with dozens to hundreds of localizations per μm^2^ (Figure 1f, zoom). In the MCF10A cells, we did not observe higher-order structural features of Satb1 (such as tendrils or droplets), and there were no apparent sub-nuclear spatial preferences (Figure 1f, Supplementary Figure 1c). The substantial (100-fold) spatial variation of Satb1 localizations could reflect heterogeneity in the distribution of Satb1 binding sites throughout the genome and/or slow (but cooperative) binding to random subsets of sites.

To investigate the origins of Satb1’s binding heterogeneity we studied μm^2^ sized chromatin regions and asked whether Satb1 occupied some regions more frequently than others. When we summed the intensity data from the entire experiment, we noted bright spots that stood out from their local background (Figure 1g). These regions were characterized by the repeated arrival, capture, and departure of Satb1 molecules (Figure 1h-i). Next, we employed several deep sequencing tools to define the genomic sequence, nucleosome occupancy, and torsional state of the genomic DNA in regions preferred by Satb1.

### Genome mapping reveals a dominant Satb1 consensus motif across multiple cell types

To explore the genome-wide distribution of Satb1, we performed chromatin immunoprecipitation followed by deep sequencing (ChIP-seq) in VL3-3M2 and MCF10A cells. The ChIP-seq data were highly replicable (Pearson correlation coefficient for ChIP-seq duplicates > 0.995, Supplementary Figure 2). Figure 2a, b shows a heat map centered on the Satb1 binding peaks and sorted by peak intensities in MCF10A cells. A genome-wide scan for Satb1 targets revealed ∼22,000 non-redundant binding sites in MCF10A and ∼4000 sites in VL3-3M2, mostly located in intergenic and intronic regions (Figure 2d, Supplementary Figure 3a, d). To validate whether Satb1 binding sites preferentially localized to A/T rich regions, we calculated the A/T percentage in the same symmetric 4kb window as used for calculating ChIP-seq intensities. As expected, the A/T percentage heat-map was strongly correlated with ChIP-seq signal (Figure 2b, c; Supplementary Figure 3b, c). To identify sequence motifs bound by Satb1, we performed *de novo* motif discovery using MEME^42^ on a 150bp DNA sequence centered on each ChIP-seq peak. For MCF10A, 91% of all binding sites were satisfied by a single consensus motif (Figure 2e) whereas in VL3-3M2 two major motifs covered 100% (Supplementary Figure 3e) and 22% (Supplementary Figure 3f) of all binding sites, respectively. To test if the A/T rich inverted palindromes predicted by SELEX^32^ constituted a subset of the major motif class in MCF10A (Figure 2e), we searched for the two most common motifs predicted by SELEX^32^. We found only 192 occurrences of TATTA**G**TAATAA and 64 occurrences of TATTA**G**TAATAC out of 22829 peaks, underlining the limitations of previous in vitro SELEX binding data in determining Satb1 consensus .

**Figure 2.**
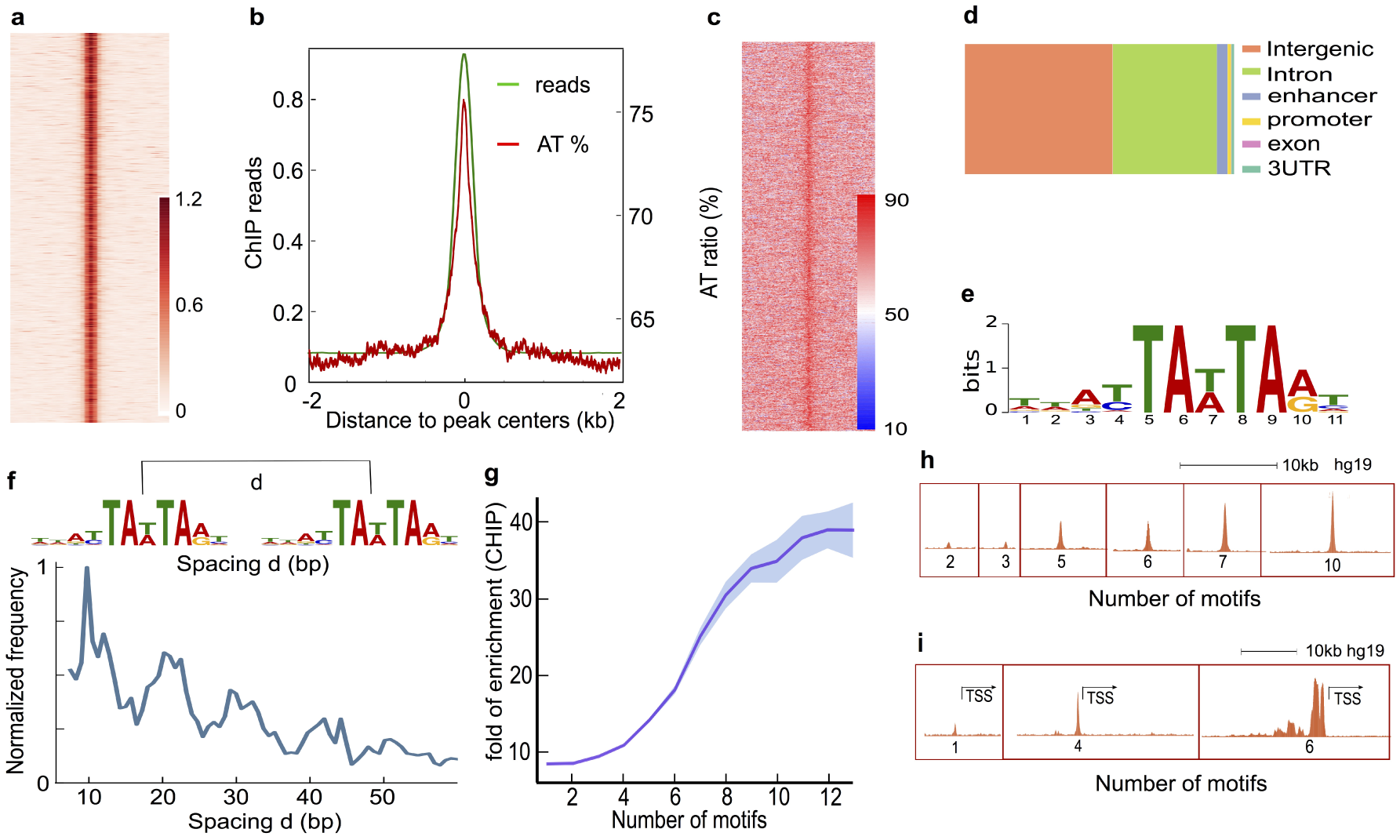
**Genome wide binding of Satb1 reveals a major consensus motif, clustered distribution of binding sites and cooperative binding. a**. Heatmap of ChIP-seq signals (read pileups) along a 4kb window centered on binding peaks. **b**. Overlay of average ChIP-seq signals (green) and AT percentage (red) along the same 4Kb window as in (a). **c**. Heatmap of AT percentage along the 4kb window as in (a). **d**. Relative abundance of Satb1 binding sites in different genomic categories. **e**. Identification of a Satb1 consensus motif in MCF10A using MEME. **f**. Frequency distribution of the spacing between Satb1 bound motif pairs in a 400bp window, showing ∼ 10bp periodicity. **g**. Satb1 ChIP-seq shows cooperative binding (mean ChIP-seq signal versus number of motifs per binding site reveals a sigmoidal binding curve with a Hill coefficient of 5.3. **h, i**. Representative examples of ChIP-seq signal strength classified by motif density per binding site in gene body **(h)** and proximal to TSS **(i)**.

### Periodicity and dense motif clustering characterize *in vivo* Satb1 binding regions

The most visually prominent feature of Satb1’s distribution in MCF10A cells are the chromatin interaction hotspots. Do these hotspots correspond to regions of the chromatin that are highly enriched for Satb1 binding motifs? To gauge whether Satb1 binding sites are spatially clustered within the genome, we first detected all instances with two Satb1 motifs within a 400 bp window centered on a ChIP-seq peak (Figure 2f). Further analysis revealed that these dual sites tended to be separated by ∼10bps and that some regions had four or more such sites with ∼10 bp spacing (Figure 2f). A similar analysis for 8 other highly abundant transcription factors including several pioneer factors showed no such periodicity in motif spacing (Supplementary Figure 4a), suggesting that ∼10 bp inter-motif spacing is a unique structural constraint recognized by Satb1. Satb1 has been shown to bind well to BUR repeats *in vitro* and it has been speculated that the dimeric/tetrameric organization of Satb1 helps to stabilize its interaction with DNA through a ‘monkey bar’ geometry over long^43^ as well as short^34^ distances.

To assess whether motif density affected binding strength we plotted peak intensities against motif density in a 400 bp window centered on the ChIP-seq peak. Remarkably, the ChIP-seq signal strength increased cooperatively with motif density, with a notable Hill coefficient of 5.3 (Figure 2g). A similar analysis of motif density versus ChIP-seq signal did not reveal cooperative binding for eight other transcription factors (Supplementary Figure 4b). Figure 2h shows several examples of signal strengths at different motif densities. For Satb1, higher motif density correlated with higher signal strength, including regions surrounding the transcriptional start sites (TSS). These findings indicate that motif density is an essential criterion in defining Satb1 site selectivity through cooperative recruitment of Satb1 molecules.

### Preferential binding of Satb1 to dense nucleosome regions is akin to pioneer factor behavior

A genome-wide search of the Satb1 sequence consensus in MCF10A cells revealed ∼2.6 million perfect sequence matches. Yet *in vivo*, Satb1 only binds a tiny fraction (∼0.8%) of these potential targets. We asked whether nucleosomes act as barriers to Satb1 binding since the ENCODE consortium has shown that most transcription factors bind to highly accessible (nucleosome-free) DNA regions.^11,12^ We used ATAC-seq^40^ to relate Satb1 binding sites to genome accessibility in MCF10A cells. For both native MCF10A cells and engineered cells expressing exogenous Satb1, the ATAC-seq fragment length distribution was enriched in nucleosome-free regions and within a dominant mono-nucleosome peak (Supplementary Figure 5a). As expected, the general ATAC signal was highly enriched near transcriptional start sites (TSS, Supplementary Figure 5b, c). Also as expected, positive control data for CTCF binding (which prefers nucleosome-free regions) strongly overlapped with high ATAC signals (Figure 3a). In sharp contrast, the bulk of the Satb1 target sites (∼95%) fell in transposase-inaccessible regions (Figure 3b). Compared to Satb1 targets in transposase-accessible regions, a significantly higher fraction of Satb1 bound sites in transposase-inaccessible regions localized to promoters and enhancers (Figure 3b). For Satb1 binding sites located in both inaccessible and accessible chromatin, the ATAC signal was visibly depleted (Figure 3c, d), suggesting that Satb1 could be binding directly to nucleosomes.

**Figure 3.**
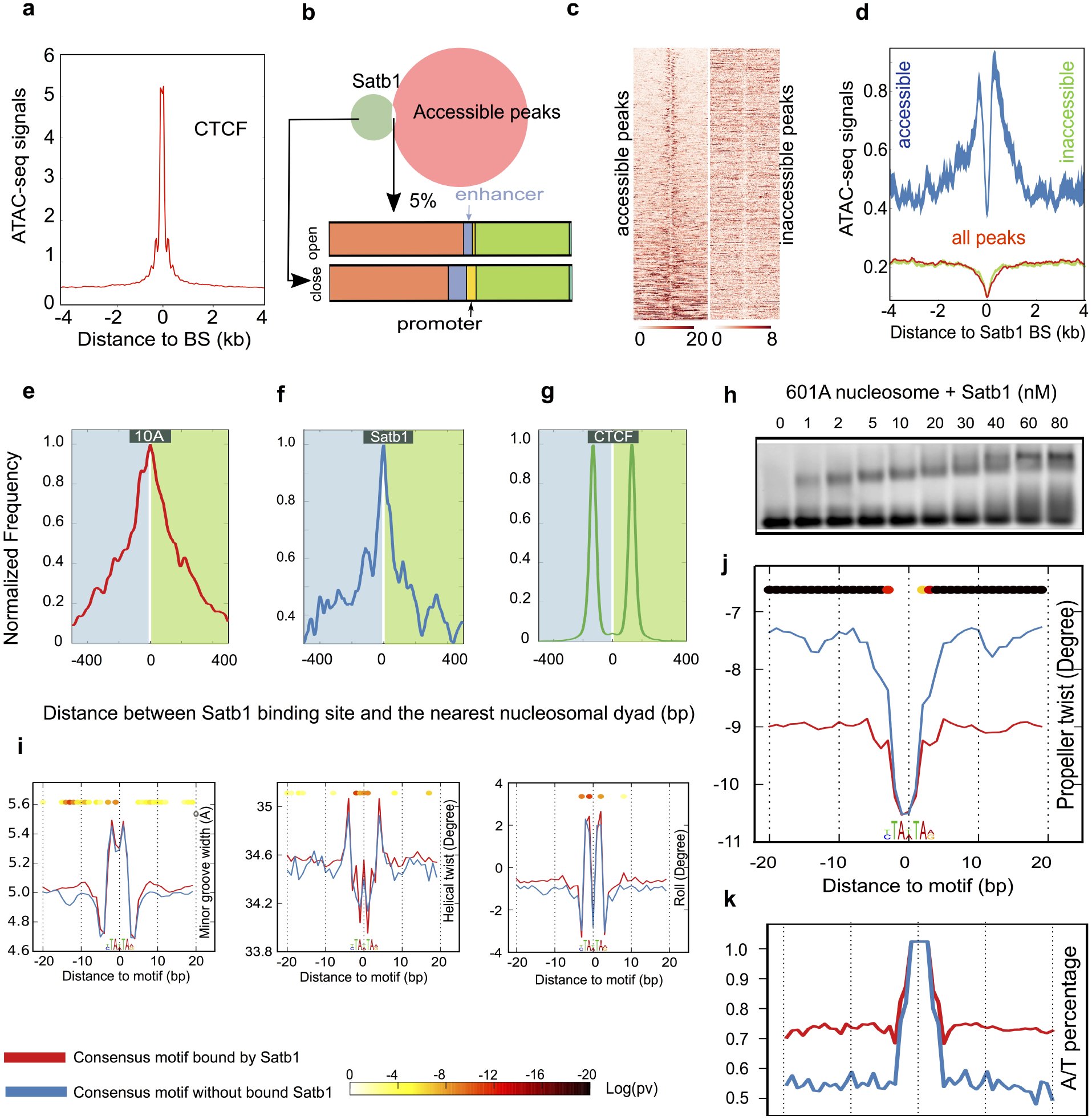
**Satb1 binds mostly to nucleosome dense regions and to sites with flanks characterized by enhanced negative propeller twist**. **a**. The average of ATAC-seq tagmentation signal from an 8kb genomic window centered on CTCF binding sites in transposase accessible and inaccessible regions of the genome. **b**. Genome annotation for accessible (∼ 5% of all binding sites) and inaccessible Satb1 binding sites (∼95% of all binding sites). **c**. The ATAC-seq tagmentation signal from an 8kb genomic window centered on Satb1 binding sites in transposase accessible and inaccessible regions of the genome. **d**. The average of tagmentation signals at Satb1 binding sites in accessible and inaccessible regions. **e**, **f.** Distribution of distances from nucleosomal dyad to Satb1 binding centers in native MCF-10A cells which lack Satb1 **(e)** and cells expressing exogenous Satb1 **(f)**. **g**. Distribution of distances from nucleosomal dyad to CTCF binding centers in MCF-10A cells. **h.** 601A nucleosome-EMSA shows that Satb1 binds efficiently to nucleosome-core embedded consensus motif resulting in 3 distinctly shifted populations, each increment presumably representing Satb1 binding to one of the sites incorporated in SH3, SH4 and SH6. **I, j**. Line plots showing average of MGW, HelT, Roll **(i)** and ProT **(j)** values across 40 bp windows centered on consensus motifs bound by FL Satb1 in nucleosome free regions. **k**. Line plots showing average percent A/T content across 40 bp windows across the same 40 bp windows as (i,j).

### Satb1 binds to nucleosomal core sequences *in vivo* and *in vitro*

The capacity to bind nucleosome-dense regions is a hallmark of pioneer factors.^44^ Satb1 has been shown to have pioneer activity in Treg cells.^27^ Satb1’s preference for inaccessible chromatin made it inherently difficult to determine nucleosome positions at target sites due to low ATAC signal (Figure 3b-d). We enriched sequencing reads that could be mapped to less accessible regions by performing ATAC-seq on Satb1 ChIP-enriched chromatin, and by removing reads shorter than one nucleosome length (Materials and Methods). This approach markedly increased the number of binding sites for which we could map nucleosome position.^45^ The distribution of distances between the Satb1 binding center to the nearest nucleosome dyad showed that the bulk of Satb1 binding sites reside within the nucleosomal core (Figure 3e, f). This pattern is opposite to CTCF, which binds to inter-nucleosomal linker regions (Figure 3g).^40^ It has been suggested that transient unwrapping of core nucleosomal DNA allows pioneer factors to bind nucleosomes. To test whether Satb1 can bind nucleosomes directly by accessing target sites located in core nucleosomal sequences, we generated a modified version of the strong nucleosomal positioning sequence 601^46^, where minor-groove outer facing nucleotides at three different superhelical locations (SHL3, SHL4, SHL6) were modified to generate three Satb1 consensus sites (601A, Materials and Methods). Recombinant Satb1 robustly bond to 601A nucleosomes (Figure 3h), confirming the genomics data and establishing that Satb1 is capable of binding target sites inside the nucleosomal core sequence even in the absence of remodelers. While these findings strongly support Satb1’s candidacy as a pioneer factor, they do not explain Satb1’s selective binding to a small subset of putative target sites.

### Enhanced negative propeller twist in flanking regions correlates with increased Satb1 binding

To assess whether the ‘shape’ of flanking sequences affects Satb1 selectivity, we compared the occupied to the non-occupied consensus Satb1 motifs. For each motif, we retrieved 100bps on either side of the motif and evaluated its DNA shape (Materials and Methods).^47^ The sequences were further categorized into nucleosome-free regions (based on nucleosome occupancy analysis, Materials and Methods) and accessible or inaccessible chromatin (based on ATAC seq). The minor groove width (MGW), helical twist (HelT), and Roll varied only slightly with Satb1 occupancy, but bound motifs had higher negative propeller twist (PT) in their flanking sequences (Figure 3i, j). Propeller twist may be a proxy for DNA flexibility.^48^ All three sequence categories followed this pattern (Supplementary Figure 6a, b) with the signal being most robust in the nucleosome-free regions (Figure 3i, j). Because propeller twist strongly depends on the AT-content,^48^ we compared this parameter for the Satb1-bound and unbound targets. Satb1 bound sites showed higher A/T content in their motif-flanks (75%) than the unbound ones (67%) (Figure 3k, Supplementary Figure 6c). This suggests that A/T content and Propeller twist of motif flanks are important determinants of Satb1 site selectivity.

### Satb1 stabilizes BURs against unwinding under torsional stress

Classic Satb1 target sites such as the immunoglobulin heavy chain (IgH) enhancer region have been shown to melt over extended regions under torsional stress.^31,29^ Torsion can be generated by replicating and transcribing polymerases, nucleosome assembly, and remodeling, as well as chromatin condensation by ATP dependent condensins.^49,14^ Topoisomerases maintain a torsionally relaxed genome by relieving torsional strain in chromatin that can build up in the wake of transcription and chromatin remodeling.^14^ To trap the genome in a torsionally strained state we inhibited topoisomerase I (Topo I) and topoisomerase II (Topo II) in MCF10A cells for a short time with Camptothecin (CPT) and ICRF-193, respectively, and we then carried out ChIP-seq of Satb1 bound chromatin.

Upon treatment with ICRF-193, there was a marked increase in binding for all binding sites (Figure 4a). Camptothecin did not affect binding to low strength sites but increased binding to high strength sites (Figure 4a). This increase in binding strength is further evident in Figure 4b, which shows that Satb1 binds more cooperatively upon ICRF-193 as well as CPT treatment (Figure 4c). To delineate the torsional state of Satb1 binding sites we carried out TMP-seq^15^ on WT MCF10A cells and MCF10A cells stably expressing Satb1-eGFP. We also collected TMP-seq data from both of these cell types upon treatment with the topoisomerase inhibitors CPT and ICRF-193. Psoralen-crosslinking efficiency at Satb1 binding sites in native MCF10A cells diminished markedly upon CPT treatment and to a much greater extent upon ICRF-193 treatment (Figure 4d). Upon expression of exogenous Satb1, the stark deficiency in Psoralen-crosslinking at Satb1 binding sites seen in native MCF-10A cells treated with CPT and ICRF-193 was abrogated (Figure 4e). This suggests that Satb1 prefers torsionally stressed DNA and stabilizes binding sites against helix destabilization.

**Figure 4.**
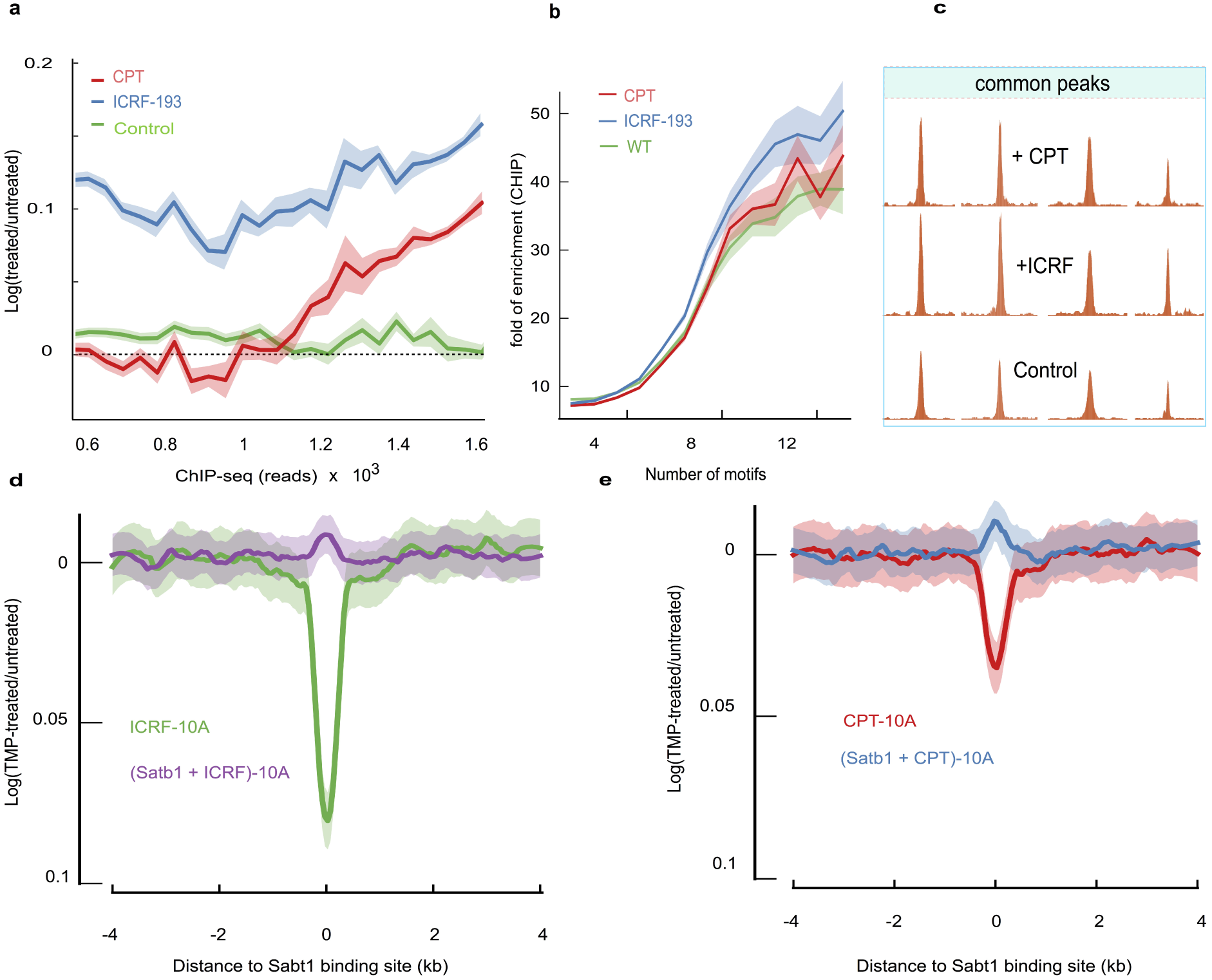
**Satb1 shows improved binding under enhanced negative torsional stress and stabilizes BURs upon binding. a**. Satb1 ChIP-seq signal ratio of “MCF-10A+Satb1” cells treated with Topo I or II inhibitors, ICRF-193 (blue) and CPT (red) respectively over untreated “MCF-10A+Satb1” cells, plotted against increasing raw ChIP-seq reads. The green line shows the ratio between two wild-type replicates. The logarithm plot of ratio shows increased Satb1 binding upon treatment. **b**. Plot of ChIP-seq signal of ICRF-193 (blue), CPT (red) treated “MCF-10A+Satb1” cells vs. motif density calculated from a 400bp window centered on Satb1 binding sites. **c**. Few representative examples of ChIP-seq peak intensity before treatment (lower panel) and after treatment with ICRF-193 (middle panel) and CPT (upper panel). **d, e**. TMP-seq was performed on native MCF10A cells and cells expressing exogenous Satb1 in presence and absence of CPT and ICRF-193 respectively. Final TMP-seq signal was derived after correction for free genomic DNA-sequence bias at 50 bp resolution across a 4kb window centered on Satb1 binding sites. Ratio of TMP-seq signals in “MCF-10A+Satb1” cells treated with CPT (**d**) or ICRF-193 (**e**) over untreated “MCF-10A+Satb1” cells.

### Protein Domain determinants of Satb1 binding affinity and specificity

A mechanistic explanation of the context sensitivity of how Satb1 interacts with the chromatin requires an understanding of how a single protein can integrate multiple selection criteria. For example, consider an additive model where Satb1’s three (putative) DNA interaction domains (CUT1, CUT2, and HD) all contribute to affinity and specificity; alternatively, affinity and specificity could arise out of the modular organization of Satb1 where some domains could provide affinity and others would provide specificity. Since several HD family transcription factors have been shown to bind to sequence environments with enhanced negative propeller twist^50^, it is plausible that the Satb1 HD might play a role in conferring specificity through the recognition of specific DNA shape parameters. To address these questions, we compared full-length Satb1 (‘**FL**’) to three domain truncation mutants, one with the N terminal dimerization domain but no DNA interaction domains (‘**N**’), one with the N terminus and CUT1 (‘**N-C1**’), and one lacking the HD (‘**∆HD**’) (Figure 5a). We used a combination of imaging and genomic techniques to evaluate the ability of these different domain mutants to (a) minimally bind chromatin, (b) stably bind chromatin, and (c) specifically bind subsets of Satb1 motifs.

**Figure 5.**
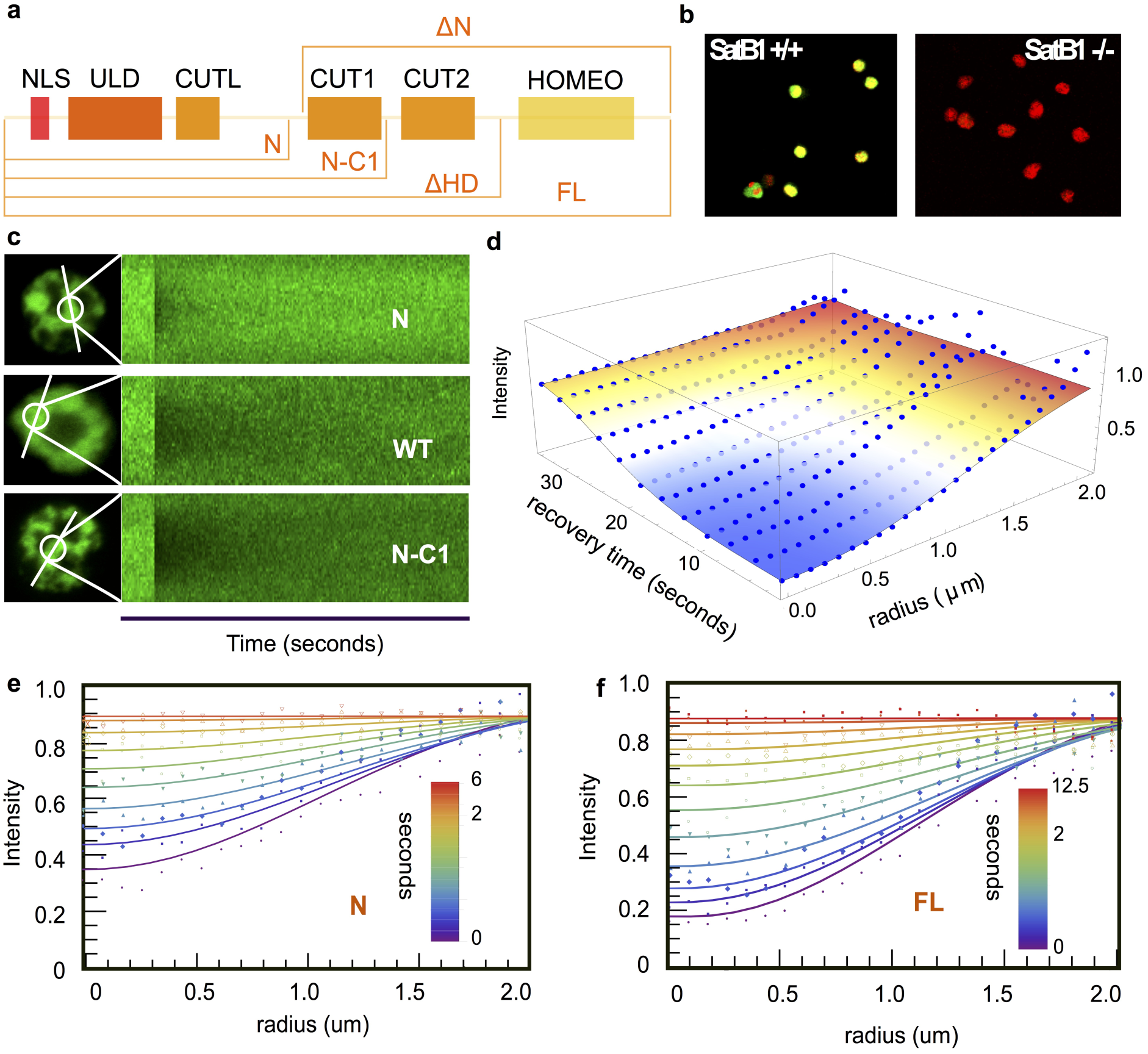
**Spatiotemporal FRAP analysis of Satb1 dynamics in the nucleus. a**. A schematic representation of the domain organization of Satb1 and the various domain constructs which were generated to study Satb1 dynamics. **b**. Satb1 knockout lines were generated by CRISPR/Cas9 strategy. Immunofluorescence image of WT Vl3 3M2 cells and Vl3 3M2 cells where the native Satb1 has been knocked out (Satb1 green and DNA in red) **c**. Examples of time vs fluorescence intensity carpets of high resolution spatiotemporal maps of fluorescence bleaching and recovery along scan lines bisecting a bleach spot in Satb1^-^/^-^ cells expressing exogenous eGFP fusions of **FL** Satb1 and different domain constructs. **d**. The best-fit surface to a spatiotemporal FRAP data. **e**, **f**. The best fits of representative recovery profiles for **N** (**e**) and **FL** (**f**) are displayed at select times, with the color bar representing time in seconds.

We generated VL3-3M2 cell lines where the native Satb1 was knocked out using CRISPR/CAS9 (Materials and Methods, Supplementary Figure 7a, b; Figure 5b) and was replaced with an inducible version of a full-length Satb1 or a domain truncation mutant. Spatiotemporal FRAP^51,35^ was used to determine the protein’s affinities to chromatin (Figure 5c-f, Materials and Methods). The 2D spatiotemporal profiles were fit with a reaction-diffusion model^52^ (Figure 5e-f, Materials and Methods, Table1). Notably, all constructs containing CUT1 were able to stably bind chromatin with similar dynamics, suggesting that CUT1 is the primary determinant for stable chromatin interaction. Consistent with earlier *in vitro* findings^43^, **N** retained minimal binding to chromatin (Figure 5c, e; Table 1).

**Table 1:**
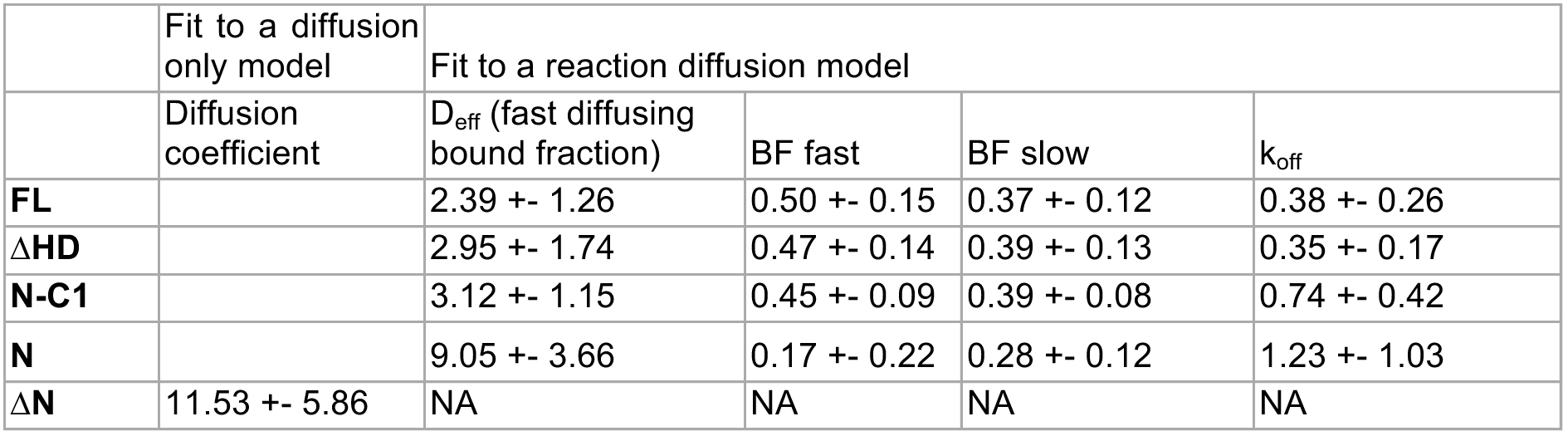
**Estimates of spatiotemporal FRAP fit parameters**

In addition to FRAP, we carried out single particle tracking of **FL** Satb1 and different domain truncations to directly visualize binding (Figure 6a, Supplementary Movies 1-4). For all four cell lines, a typical cell yielded ∼2000 localizations (Supplementary Figure 8a). The comparable number of localizations across the four conditions reflects similar expression levels and uniform imaging settings. The molecular position vs. time tracks (Figure 6b) revealed two major classes of Satb1, namely slow (and likely anchored to chromatin) and fast (and presumably diffusing through the nucleus). Some trajectories showed transitions between states (Figure 6b, ‘multistate’). On a time-projection of the individual frames, the long-lasting tracks were visible as contiguous signals (Figure 6c, d).

**Figure 6.**
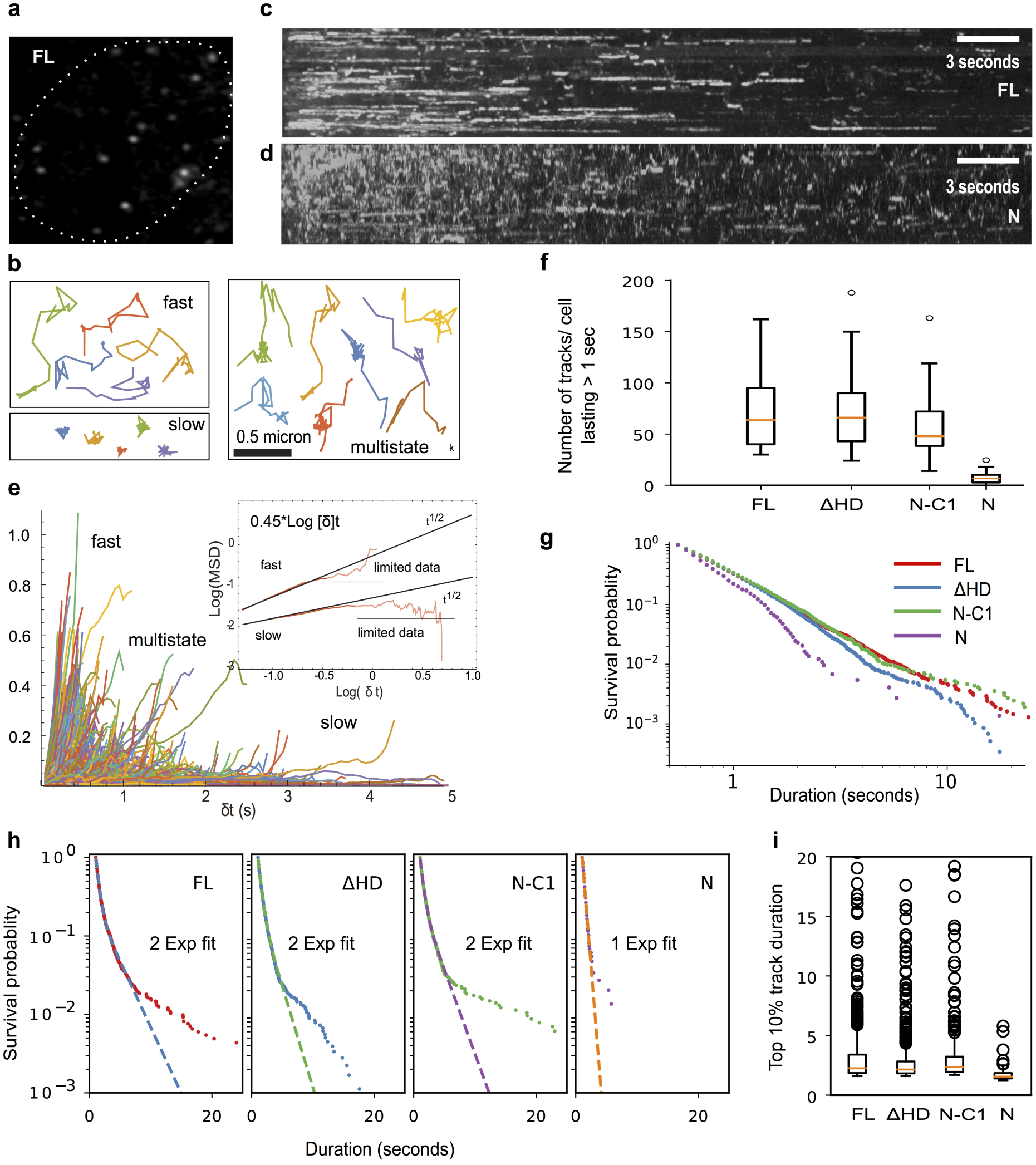
**Single molecule tracking analysis of Satb1 in the nucleus. a**. A Hilo TIRF image of an MCF-10A cell with leaky expression of Satb1-eGFP showing diffraction limited spots of single Satb1 molecules. **b**. Examples of FL tracks at 20 Hz. The maximum inter-frame displacement was set to 0.25 um which is the maximum step size that we observed for nucleosome embedded histone H2B (see Supplementary Figure 8b-d). Displacements are classified into fast, slow, and multistate categories (See Methods). The latter trajectories consist primarily of molecules that are initially strongly confined, but then exhibit larger displacements. **c**, **d**. Time projection stacks reveal stably bound molecules as continuous stretches of signal. FL (**c**) time projection has more instances of contiguous signal than the dimerization unit **N** (**d**). **e**. Evidence of two states and subdiffusion. **f**. Long tracks are enriched with N-C1, ∆HD, and FL. Box whisker plots showing distribution of number of tracks per cell that last more than 1 second (20 consecutive frames). Median values for **D** (6), **N-C1** (48), **∆HD** (66), **FL** (64). **g**. Survival plot of track durations for **FL** and the different domain truncations show that the Homeo domain is dispensable but at least one CUT domain is essential for longer residence times. **h**. For **FL**, **N-C1** and **∆HD** the survival distributions are fit better with two exponentials than one except for **N**, which exhibits mono-exponential distribution of bound molecules. **i**. Box-whisker plot representing the distribution of track durations that constitute the top 10% of residence times for the different Satb1 constructs.

Since ensemble averaging over a heterogeneous population obscures motion characteristics (Figure 6e), we grouped trajectories in slow and fast fractions based on their initial displacements (Figure 6e inset). The ‘fast’ and ‘slow’ MSD data for small time intervals were well fit by lines of constant slope of ∼1.02 and ∼0.45, respectively (Figure 6e inset). Single locus tracking experiments in Yeast^53^ and other eukaryotes^54^ have found values of ∼0.5 for the exponent of locus diffusion^53^, similar to our findings of ∼0.45 (Figure 6e inset). This suggests that when Satb1 is bound to chromatin, it exhibits the Rouse dynamics^53^ of the slowly undulating chromatin polymer.

The effective diffusion coefficient for the ‘slow’ fraction of around 0.04 μm^2^/s is consistent with previous measurements for chromatin fiducials.^53–54,55^ The diffusion coefficient estimated for the ‘fast’ faction, around 0.15 μm^2^/s, falls on the lower bound of what has been measured for freely diffusing transcription factors (typical values of 0.5 to 5 μm^2^/s). For each construct, we determined the distribution of dwell times (Figure 6f-i). Consistent with FRAP data, **N** showed very few tracks with >1 second lifetime (Figure 6f), suggesting that **N** can interact with chromatin, either directly or indirectly but does not engage stably (Figure 6f-g; Supplementary Movie 4). The other constructs exhibited a significantly higher fraction of long-lived tracks (Figure 6f-g, Supplementary Movies 1-3). For fast image acquisition rates, it has been shown that the slow fraction is dominated by outliers.^37^ A box plot of the top 10% of longest surviving tracks showed that there are many more statistically classified outliers for **FL**, **N-C1**, and **∆HD** (lasting up to 25 seconds) than for **N** (Figure 6i). Furthermore, the survival distribution of tracks lasting greater than 0.5 seconds was fit better with a biexponential function^37^ for **N-C1** and **∆HD,** unlike **N** which showed a mono-exponential distribution (Figure 6h).

To assess if HD imparts specificity, we carried out ChIP-seq on MCF10A cells inducibly expressing eGFP fusions of all four constructs. The ChIP-seq data replicates were highly correlated for all the domain constructs (Supplementary Figure 9a). In line with the single molecule data, all three constructs containing CUT1 engaged stably with chromatin (7.4 to 11.1-fold enrichment over the background, Figure 7a-b, Supplementary Figure 9b). The **N** construct pulled down comparable amounts of DNA as the other mutants and had no consistent binding pattern across the genome (Figure 7a). Interestingly, **FL** bound to significantly *fewer* genomic sites (∼22,000) than **N-C1** and **∆HD** (∼40,000 and 47,000 respectively, Figure 7c, d). Motif analysis using MEME^42^ revealed that all three constructs bound essentially the same core sequence motif (Supplementary figure 9c) and that the majority of these sites were in intronic and intergenic regions of the genome (Supplementary Figure 9d). However, the subset of sites that were exclusively bound by **N-C1** and **∆HD** had a lower A/T percentage than **FL** (Figure 6e-g, Supplementary Figure 10a-c), lower motif density, and showed no signs of cooperativity (Figure 6h inset). DNA shape calculations revealed that these sites were characterized by a more positive propeller twist angle and lesser A/T content in their immediate flanks. This was true for both nucleosome-free regions (Figure 6i, j) and inaccessible regions (Supplementary Figure 11a, b). In summary, these data indicate that CUT1 is sufficient for high-affinity binding whereas HD primarily aids in specificity through shape readout of motif flanks.

**Figure 7.**
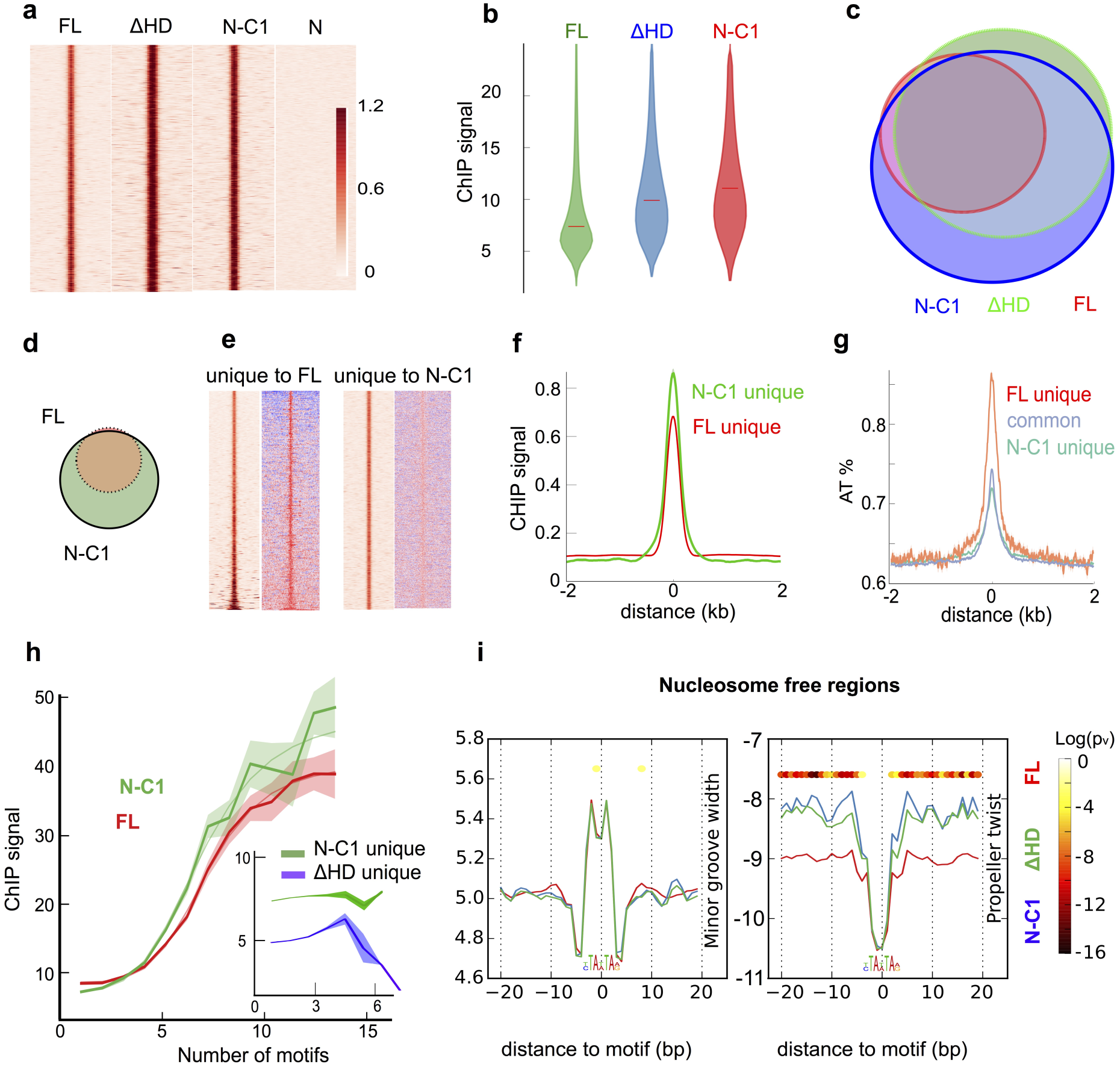
**Homeo domain is dispensable for high affinity binding but increases site selection stringency through DNA shape recognition. a**. Heatmap of ChIP-seq signals (read pileups) along a 4kb window centered on binding peaks of different Satb1 domain constructs. Both **N-C1** and **∆HD** show higher signal enrichment at binding sites than **FL** Satb1. The dimerization domain **N** on its own shows no consistent signal enrichment. **b**. Violin plots showing the distribution of the peak signal strengths for 3 different Satb1 constructs (Median peak strengths are shown as horizontal lines). **c**. Venn diagram showing binding site overlap between all 3 domain constructs. **d**. Venn diagram of **FL** vs. **N-C1** binding sites. **N-C1** binds nearly all (∼97%) of all **FL** binding sites and more than as many sites exclusively. **e.** Heatmap of ChIP-seq signals (read pileups) (left column) and AT percentage (right column) of binding sites unique to FL Satb1 and **NC1**. **f**, **g**. Average plot of ChIP-seq signals (**f**) and AT percentage (**g**) along a 4kb window across the binding sites that are exclusively bound by FL Satb1 or **N-C1**. **h**. **N-C1** and **∆HD** bind cooperatively to sites that are common with **FL** Satb1. However, sites exclusively bound by **N-C1** and **N-C1-2** show no such cooperative pattern (inset). **i**. Distinct DNA-shape features help define **FL** Satb1 and **N-C1** binding. Line plots showing average of MGW and ProT values across 40 bp windows centered on consensus sites bound by **FL** Satb1 and sites bound exclusively by **N-C1** and **∆HD** in nucleosome free regions.

### Conclusion

Our data provide the first comprehensive quantitative picture of how Satb1 uses multiple identifiers to selectively target a small cohort of potential binding sites in the genome. SMT and spatiotemporal FRAP revealed that the CUT1 domain is sufficient to ensure prolonged interactions with the genome. By complementing imaging with deep sequencing, we have found that while HD is dispensable for prolonged interactions in SMT experiments, it plays a crucial role in assigning specificity by selecting targets surrounded by sequences with more negative propeller twist and higher A/T percentage. Further, Satb1 shows cooperative binding to genomic regions with high binding site density, a feature that is also visible in dynamic super-resolution images as chromatin interaction hotspots. While long tracts of poly(dA:dT) are unfavorable to nucleosome formation,^56^ bendable di-nucleotides (AT, TA) frequently occur periodically at helix/histone interaction sites along the length of a nucleosome.^56^ Interestingly, Satb1 binding sites also show ∼10 bp periodic spacing and Satb1 preferentially targets closed chromatin by directly accessing nucleosome embedded motifs *in vivo* and *in vitro*. Indeed, enhancement in Satb1 binding efficiency upon treatment with CPT and ICRF-193 is consistent with a nucleosome mediated cooperativity model,^57^ as treatment with topoisomerase inhibitors would destabilize nucleosomes, facilitating partial unwrapping. Torsional stress sensing and preferential binding to nucleosome embedded sites may constitute the effector and response modules of Satb1’s binding mechanism. It has been suggested that recognition of partial motifs on one face of nucleosomal DNA is the discerning feature of a pioneer factor.^44^ Further studies are needed to delineate the Satb1-nucleosme binding interface at high resolution and to temporally resolve the changes in chromatin architecture upon binding of Satb1 to silent chromatin.

## Materials and Methods

### CRISPR

We knocked out native Satb1 in Vl3 3M2 with CRISPR-mediated homogeneous recombination technique. Source vectors of guide-tracer DNA and Cas9 was obtained from Doudna lab at Berkeley and Zhang lab at MIT, through Addgene. Briefly, we first designed a CRISPR guide vector with double BsaI sites that is able to insert target oligo efficiently using golden gate method. This vector was constructed using SLIC.^58^ The donor plasmid was built based on pmCherry-C1 vector where the DNA of two homologous arms flanking the sequence of interest was inserted. Here mCherry was used a negative selection marker as it would be removed if proper homologous recombination and integration took place. The HR cassette consisted of a 5’ end and 3’ end homologous arms of 800-1000bp (PCRed directly from Vl3 3M2 genomic DNA) flanking the loci of interest (for KO: puromycin-BGH polyA, and for HR eGFP fusion at 3’ end: flexible linker-eGFP-P2A-puromycin). Surveyor assay was performed for each guide sequence according to Sanjana et al. 2012.^59^ HR donor plasmid, guide DNA plasmid and Cas9 cDNA plasmid were electropolated into Vl3-3M2 cells using Neon system (invitrogen). Double nickase version of cas9 was used to enhance the editing specificity.^60^ The cells were then selected with puromycin after 3 days of culture and sorted into 96-well for clonal selection. Cells per well was immuno-stained to obtain the final clonal Vl3 3M2 cells with double-allele knock-out. To tag the native Satb1 with eGFP in Vl3 3M2 cells, we used the same technique to insert an eGFP-P2A-puromycin sequence cassette replacing the stop codon by homologous recombination.

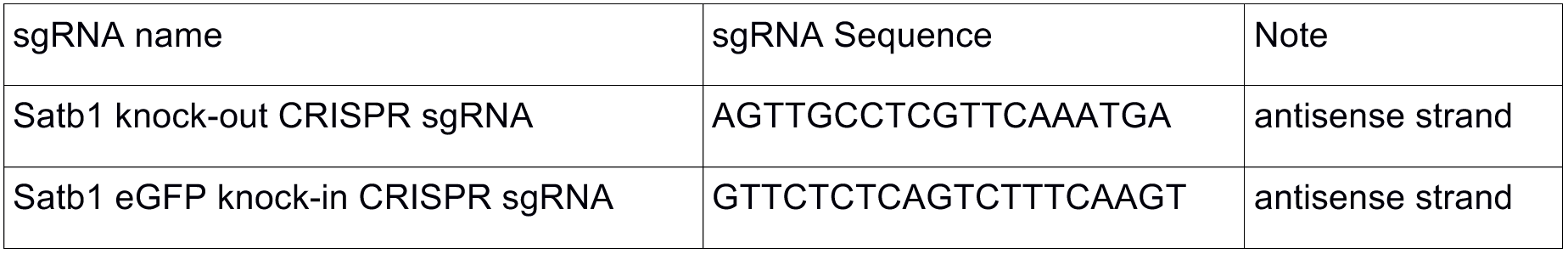

### PLASMIDS and cell lines

FL Satb1 and the different domain deletion mutants were generated as C terminal eGFP fusions in a pLenti based vector with cumate switch promoter and stably integrated into VL3 3M2 Satb1 ^-^/^-^ or MCF-10A cell lines. Stable cell lines were generated by lentiviral transduction. Cells were then selected for appropriate resistance marker to achieve stable integration and sorted by FACS.

### Protein purification

Recombinant pETDuet plasmids were transformed into SoluBL21 *E. coli* (Genlantis, cat. C700200). *E. coli* were grown in M9 minimal media (according to manufacturer’s protocol) at 16C, and protein expression was induced by IPTG. Unless otherwise stated, all subsequent steps were done at 4 C. Cells were spun down and lysed prior to protein isolation by standard His Nickel-NTA (Invitrogen, cat. R90101) column chromatography. The Nickel-NTA isolated fraction was further purified using Superose 6 Increase column 10/300 GL (GE Healthcare) size-exclusion chromatography. The choice of column provided the appropriate fractionation range and resolution such that the expected protein tetramer could be purified from smaller incomplete or degraded products. Subsequent analysis of column fractions was performed by visual inspection of SDS-Page gel electrophoresis (Invitrogen, cat. NP0322) and fractions meeting expected size and purity were pooled and concentrated (50 mM TrisHCl, 100 mM NaCl, 1mM EDTA, 1mM DTT, 10% v/v Glycerol, pH = 7.3 at 25 °C).

### Nucleosome reconstitution and EMSA

Core histone octamers were isolated from chicken erythrocytes as described previously.^61,62^ Nucleosomes were assembled using 5’ Alexa 488 labeled 601A which is derivative of the nucleosome positioning 601.^46^ The nucleotides at superhelical locations SHL3, SHL4 and SHL6 (where the minor groove is facing histones/histone core) were modified (shown below in bold) to generate Satb1 consensus sites.

5’CTATACGCGGCCGCCCTGGAGAATCCCGGTCTGCAGGCCGCTCAATTGGTCGTAGACAGCTCTAG CACCGCTTAAACGCACGTACGCGCTGTCCCCCGCGTTTTAACCGCCAAGGT**TAATA**ATCCT**TAATA**A CCAGGCACGTGTCAT**TAATA**AACATCCTGTGCATGTGGATCCGCACTC3’

Reconstitution was performed as described earlier.^62^ For Electrophoretic Mobility Shift Assays 6 nM nucleosomes were incubated with increasing loading of Satb1in total volume of 25 ul in STE buffer (80 mM NaCl, 20 mM Tris-HCl, pH 8.0, 0.01% NP-40, 1 mM DTT). Resulting complexes were separated on 1% type IV (Sigma cat. #A3643) agarose gel electrophoresis in 0.5xTBE buffer at constant 100 V and visualized on Typhoon fluoroimager.

### Image correlation spectroscopy

Image correlation spectroscopy was conducted according to Cardarelli *et al* 2010.^36^ Zeiss LSM 700 laser scanning microscope was used for image acquisition using 63x oil immersion NA 1.40 objective. Data were processed by the SimFCS software developed at the Laboratory for Fluorescence Dynamics.

### Single molecule imaging

All live cell single molecule imaging was done using HiLo Total internal reflection microscopy (HiLo-TIRFM)^41^, on an Olympus CellTIRF system. TIRF angles for each fiber-coupled illumination laser were controlled independently with a motor. Imaging was done through a 1.49 NA 100x objective and 1.6x Optovar magnifier and recorded on an Andor iXon Plus EMCCD camera at 20 Hz. All dichroics and filters were purchased from Semrock. All experiments were performed within a heated, CO2 controlled incubation chamber set to 37 °C and 5%, respectively with additional temperature control provided using a collar-type objective heater, also set to 37 °C.

### Single Molecule tracking and analysis

Single molecule tracking was preformed using Mathematica and python. Detectable particles in each frame were first identified by performing a Laplacian of Gaussian filter and then thresholding the filtered image based on intensity.

Then the raw image data of each identified particle was fitted with a 2D Gaussian function to obtain particle x/y position and signal amplitude. The single particle positions in each frame were then linked into single particle trajectories based on nearest-neighbor algorithm with a defined maximum per-frame jump distance. The maximum frame to frame displacement was set to 0.25 μm as this was the maximum step size that we observed for the chromatin fiducial histone H2B. Finally, a quality control step was performed where a large subset of particles and trajectories were validated by eye. Trajectories that lasted ≥10 frames (0.5 seconds) were selected for further analysis. Individual Satb1 molecules showed considerable variability in their motion. We classified displacements into fast, slow, and multistate categories based on the initial MSD (mean squared displacement) values of each trace. The first 10 Log[δt] points (corresponding to δt = 500 ms) were fit to a line (Log[MSD] = a + b*Log[δt]).

For super-resolution imaging, after localizing the fluorophores, we combined all the localization centroids for the full length Satb1 in one computed image. For each detected molecule, a normalized symmetric 2D Gaussian (with a standard deviation equal to the computed localization uncertainty, which is on order of 30 nm) was drawn, and all the Gaussians from all the localizations (collected over 35 seconds @ 20Hz) were summed to yield a super-resolution image.

### Spatiotemporal FRAP imaging and analysis

For FRAP experiments VL3 3M2 cells expressing different domain constructs were grown in RPMI media and then embedded in CyGEL Sustain (#CS20500) Biostatus for live imaging. FRAP experiments were performed on a Zeiss LSM 700 confocal microscope with a 63x/1.40 oil immersion objective. Cells were equilibrated at 37C with 5% CO2 for 20 minutes before imaging. For different cell lines, we scanned a 50 x 2-pixel rectangle (pixel size = 0.0893 μm) inside the thymocyte nucleus in euchromatic regions with homogeneous fluorescence distribution using 488nm laser with 0.6% power and PMT gain 825. The cells were imaged at a frequency of 0.0076s/frame. A circular spot with radius = 15 pix (1.34 μm) was bleached so that the center of the circle aligned with the center of the scanned rectangle. Experiments were repeated in the same cell nucleus one more time without the bleaching step to correct for photo-bleaching.^52^

The FRAP images were corrected for photo-bleaching and then normalized. Spatiotemporal fluorescence data for fitting were generated for each time point by averaging over the two pixels that constituted the height of the rectangular scan region. This resulted in a 50 x 1 pixel-wide line that represented the fluorescence along a line bisecting the bleach spot. This line was then averaged about the bleach center to produce a 25 x 1 pixel line that represented the radial distribution of the bleach spot fluorescence. These data were exponentially sampled in time and fit with a reaction diffusion model.^5235^ Three parameters: D_eff_, k_on*_ and k_off_ were extracted from the model. Data processing and model fitting were designed and implemented in *Mathematica*.

The FRAP parameters extracted from the fit have been elaborated in Stasevich *et al*.^35^ Briefly, FRAP recoveries of Satb1 variants were first fit to a pure diffusion model, which yielded the diffusion constant *D*. Of the five Satb1 truncation mutants, only ∆N was well fit by the diffusion model. The other four Satb1 mutants yielded diffusion coefficients that were too slow to be explained by pure diffusion. Therefore, their FRAP recoveries were fit with a reaction-diffusion model and yielded estimates of D_eff_ (effective diffusion constant). The remaining parameters b_f_ (bound fraction in the fast diffusing state), b_s_ (bound fraction in the slow diffusing state), k_on*_(association rate), k_off_ (dissociation rate) and K_d_ (dissociation constant) reported in Table 1 were calculated from the fit parameters as explained in Stasevich *et al*.^35^

### Genomic library preparation

**ChIP-seq** libraries were prepared according to Myers lab protocol^63^ with modifications to adapt to chromatin shearing using Covaris AFA. Briefly, the cells were fixed with 1% PFA for 5 minutes, washed twice with PBS and then quenched with glycine (125mM) for 5 minutes. Cells are lysed with cell lysis buffer (0.5% Triton-x100, 0.25% NP-40%, 50 mM HEPES, 150 mM NaCl, 1 mM EDTA, pH 7.5) for 15 min, and sheared in TE 10 mM + 0.1% SDS buffer with Covaris (Covaris Inc) S2 according to manufactures protocol (optimal condition for fragmentation in 1 ml vial was obtained at power intensity level 4 for 10 minutes).

After centrifuging the lysates to remove debris, 2-5 μg antibody (Abcam #ab290, #EPR3895) was added and incubated overnight at 4°C. Protein G magnetic beads (Invitrogen) were then added and the mixture was shaken for 2 hours before collecting using a magnetic rack. The samples were washed 6 times (thrice with low salt buffer, once with high salt buffer, once with LiCl buffer and once with TE buffer) according to Myers lab protocol. The library was generated with Ultra II DNA kit (NEB) according to manufacturer’s protocol.

**TMP-seq** libraries were prepared according to Teves and Henikoff (2014)^15^ with some modifications: Cells were incubated with Trimethylpsoralen (TMP) (Sigma-Aldrich T6137) at a final concentration of 2 μg/mL for 10 min in the dark. Plates of cells were then exposed to 3 kJ/m^2^ of 365-nm light (Fotodyne UV Transilluminator 3-3000 with 15-W bulbs). Cells were washed with PBS 1x, scraped from the culture dish and then spun down. The pellet was then resuspended (buffer containing PBS 1X, 0.2% TritonX100 and RNaseA) and incubated at 37°C for 30 minutes. Cells were then spun down and resuspended in TE (Tris 10mM, 1mM EDTA) with 0.5% SDS and proteinase K and incubated at 50°C for 1 hr. Chromatin was then sheared with Covaris to get 200-500 bp fragments and subsequently cleaned with columns (Fisher GeneJET kit).

Cross-linked fragments were enriched by repeated rounds of denaturation and Exonuclease I (Exo I) digestion as described in Teves and Henikoff (2014).^15^ 3 μg of sonicated DNA, diluted to 250 μL, was boiled in a water bath for 10 min and incubated in ice water for 2 min. To each sample, 30 μL of 10× Exo I buffer and 10 μL of Exo I were added, and digestion was allowed to proceed for 2 h at 37 °C. Then samples were boiled and cooled as before, and 10 μL of Exo I was added for a second round of 1-h digestion. After cleanup, Exo I–digested DNA samples were subjected to enzymatic reactions for end polishing and ‘A’ tailing. Illumina adaptors were ligated using NEBNext Ultra library prep kit. After that, the 5′ strand was digested with 20 U of λ exonuclease (NEB) for 30 min at 37 °C. The DNA was purified with Ampure beads. The resulting 3′ strand was used as a template for ten rounds of primer extension in 1× pfu Ultra II HS buffer, 0.8 mM dNTP, 1U of pfu UltraII HS DNA polymerase (Agilent) and 40 nM of P7 extension primer. The resulting single-stranded products were purified with Ampure beads and eluted.

The purified products were then appended with ribo-G in 1× terminal deoxynucleotidyl transferase (TdT) buffer, 10 U TdT (NEB) and 0.1mM of rGTP in 37 °C for 30 min. The products were purified with Ampure beads.

The single-stranded ribotailed products were ligated to a double-stranded adaptor with CCC-overhangs (**CCC overhang oligo 1**: ACACTCTTTCCCTACACGACGCTCTTCCGATCTCCC, **CCC overhang oligo 2**: (Phosphate)AGATCGGAAGAGCGTCGTGTAGGGAAAGAGTGT) and final library was generated by PCR amplification (using NEB Q5 High-Fidelity 2x master mix) for 5-10 cycles.

ATAC-seq. 50k cells were collected and washed with ice cold PBS and then further washed with RSB buffer (10mM Tris, 1mM NaCl, 3mM MgCl2, pH 7.4) with 0.1% NP40, and tagmentated with 2.5ul Tn5 enzyme. The libraries were prepared as per Buenrostro *et al* (2013).^40^

### ChIP-ATAC seq

We added a ChIP step before preparing for ATAC-seq sample to enrich the reads around Satb1 binding sites for determining nucleosome positions nearby. The Satb1 ChIP was performed as described above except the Covaris shearing time was reduced to 8 minutes. After bead assisted pull down, the sample was washed once with low salt buffer, and once with RSB buffer, and resuspended in tagmentation buffer with 2 ul of Tn5 enzyme mix. The sample was then incubated at 37°C for 15min and cleaned with DNA clean up kit (Zymo Research). Final library was prepared per Buenrostro *et al*. (2013).^40^

### Genomic data analysis

Adapter sequences in sequencing reads were trimmed off with cutadapt. Quality of reads were assessed by fastqc and aligned to human genome version hg19 and mouse version mm10 with bowtie2. Typically, alignment rate was between 90-95%.

ChIP-seq binding site peak calls were made with MACS2^64^, using Poisson statistical model. Differential binding was analyzed with DESeq2 package in R^65^, using binomial statistic model. The widely used MEME suite tools were employed for motif analysis.^42^ Specifically, de novo motif discovery was performed with MEME and FIMO was used to search for occurrence of known motifs with default threshold.

Briefly, Motif analysis was performed using FIMO tools and custom python script. A 400bp window centered at each binding site was selected and fasta sequence was retrieved with bedtools. FIMO was used to search for motif occurrence and location. Then the spacing between nearest motifs for each binding site and number of motif occurrences per 400bp was calculated to correlate with ChIP-seq signals.

Other ChIP-seq data were collected from encode project, with the following access numbers:ENCFF000WJE (RelA), ENCFF002CIA (TCF3), ENCFF002CJF (Oct4), ENCFF002CHF (NFATc1), ENCFF002CJA (nanog), ENCFF231TGQ (IRF2), ENCFF388AJH (IRF1), ENCFF856FVT (FoxA1), ENCFF002DCM (CTCF). The PWM (power weight matrix) for other transcription factors and chromatin regulators were retrieved from MEME motif database.

Homer software^66^ was used for genome feature annotation, and generation of read heatmap. ChIP-seq signals were presented as either raw reads or fold enrichment over background derived from MACS2 peak calling, as noted in the text and figures. When using raw reads, signal strength was calculated by summing the ChIP-seq reads within a defined genomic window as noted in the main text and normalized against a total count of 10M reads.

For TMP-seq analysis^15^ the first 4 bases were trimmed in each read before alignment to remove the CCC of ligated adapter. After alignment, cross-linked sites (starting base of each sequencing read) were detected and expanded on either side by 20bps to smooth the distribution and converted to bigwig files for calculation. A window of 50bp was used to average the TMP-seq signals.

TMP-seq reads from purified MCF10A genomic DNA were used to correct for any sequence bias in psoralen crosslinking efficiency. Briefly, the TMP-seq reads for each experiment were first normalized to the total read count (10M reads); then those reads were divided by background reads over a 50bp sliding genomic window to calculate the true background corrected TMP-seq signals.

ATAC-seq data were analyzed according Buenrostro *et al*. (2013)^40^ and Denny *et al*. (2016)^67^. Nucleosome positions at accessible regions were called with nucleoatac tools^45^ with default parameters.

### DNA shape analysis

The PWM (position weight matrix) for Satb1 motif was constructed from the motif discovery as described above. Genome wide Satb1 motif locations were obtained using Homer package (specifically the perl script scanMotifGenomeWide.pl). The DNA sequences flanking 300bp on either side of motif were retrieved and saved in a fasta file for DNA shape analysis using the web server.^47^ All the sequences of interest and their shape parameters were categorized into 3 groups for comparison: nucleosome free region (NFR) derived from nucleosome position analysis as described above, accessible chromatin and inaccessible chromatin. When there were more than 10000 motifs per group, motifs were down sampled randomly to 5000-7000 motifs for further analysis.

When comparing the shape parameters for weak and strong binding motifs, the shape data were mapped to ChIP-seq results to obtain the corresponding ChIP-seq signals per binding site. Typically, unless otherwise mentioned, we took 600 top and bottom motifs ranked according to ChIP-seq signals to compare the shape parameters. A nonparametric Mann-Whitney U test^6^ was used to assign p-value per DNA base with typical sample size from 300-7000 sequence elements depending on group size.

## Acknowledgments

This work was partially supported by the National Institutes of Health (NIH) National Institute of General Medical Sciences (NIGMS)/National Cancer Institute (NCI) Grant GM77856, NCI Physical Sciences Oncology Center Grant U54CA143836, National Institute of Biomedical Imaging and Bioengineering (NIBIB)/4D Nucleome Roadmap Initiative 1U01EB021237 and also by NIH grant P50-HG-007735 (H.Y.C., W.J.G.). H.Y.C. is an Investigator of the Howard Hughes Medical Institute. We are grateful to Davood Norouzi for suggestions regarding DNA sequence analysis.

## Author contributions

R.P.G and J.T.L conceived the project. R.P.G and Q.S. designed research. R.P.G. and Q.S performed most experiments. M.P.R purified Satb1 protein. L.Y helped with FRAP experiments. T.N performed in vitro nucleosome binding assays. T.J.S provided guidance for performing spatiotemporal FRAP analysis. H.Y.C and W.J.G provided guidance for genomic data analysis. Q.S., R.P.G, L.Y and J.T.L analyzed data. R.P.G. and J.T.L wrote the paper. All authors helped with editing the paper.

## References

1. Hortschansky, P. et al. Deciphering the combinatorial DNA-binding code of the CCAAT-binding complex and the iron-regulatory basic region leucine zipper (bZIP) transcription factor HapX. J. Biol. Chem. (2015). doi:10.1074/jbc.M114.628677

2. Rodríguez-Martínez, J. A., Reinke, A. W., Bhimsaria, D., Keating, A. E. & Ansari, A. Z. Combinatorial bZIP dimers display complex DNA-binding specificity landscapes. Elife (2017). doi:10.7554/eLife.19272

3. Parker, S. C. J., Hansen, L., Abaan, H. O., Tullius, T. D. & Margulies, E. H. Local DNA topography correlates with functional noncoding regions of the human genome. Science (80-.). (2009). doi:10.1126/science.1169050

4. Zhang, Y., Xi, Z., Hegde, R. S., Shakked, Z. & Crothers, D. M. Predicting indirect readout effects in protein-DNA interactions. Proc. Natl. Acad. Sci. (2004). doi:10.1073/pnas.0402319101

5. Mathelier, A. et al. DNA Shape Features Improve Transcription Factor Binding Site Predictions In Vivo. Cell Syst. (2016). doi:10.1016/j.cels.2016.07.001

6. Rossi, M. J., Lai, W. K. M. & Pugh, B. F. Genome-wide determinants of sequence-specific DNA binding of general regulatory factors. Genome Res. (2018). doi:10.1101/gr.229518.117.Freely

7. Rube, H. T., Rastogi, C., Kribelbauer, J. F. & Bussemaker, H. J. A unified approach for quantifying and interpreting DNA shape readout by transcription factors. Mol. Syst. Biol. (2018). doi:10.15252/msb.20177902

8. Gordân, R. et al. Genomic Regions Flanking E-Box Binding Sites Influence DNA Binding Specificity of bHLH Transcription Factors through DNA Shape. Cell Rep. (2013). doi:10.1016/j.celrep.2013.03.014

9. Kazemian, M., Pham, H., Wolfe, S. A., Brodsky, M. H. & Sinha, S. Widespread evidence of cooperative DNA binding by transcription factors in Drosophila development. Nucleic Acids Res. (2013). doi:10.1093/nar/gkt598

10. Lelli, K. M., Slattery, M. & Mann, R. S. Disentangling the Many Layers of Eukaryotic Transcriptional Regulation. Annu. Rev. Genet. (2012). doi:10.1146/annurev-genet-110711-155437

11. Slattery, M. et al. Absence of a simple code: How transcription factors read the genome. Trends in Biochemical Sciences (2014). doi:10.1016/j.tibs.2014.07.002

12. Thurman, R. E. et al. The accessible chromatin landscape of the human genome. Nature (2012). doi:10.1038/nature11232

13. Olson, W. K., Gorin, A. A., Lu, X.-J., Hock, L. M. & Zhurkin, V. B. DNA sequence-dependent deformability deduced from protein-DNA crystal complexes. Proc. Natl. Acad. Sci. (1998). doi:10.1073/pnas.95.19.11163

14. Teves, S. S. & Henikoff, S. DNA torsion as a feedback mediator of transcription and chromatin dynamics. Nucleus (2014). doi:10.4161/nucl.29086

15. Teves, S. S. & Henikoff, S. Transcription-generated torsional stress destabilizes nucleosomes. Nat. Struct. Mol. Biol. (2014). doi:10.1038/nsmb.2723

16. Corless, S. & Gilbert, N. Effects of DNA supercoiling on chromatin architecture. Biophys Rev (2016). doi:10.1007/s12551-016-0242-6

17. Freeman, G. S., Lequieu, J. P., Hinckley, D. M., Whitmer, J. K. & De Pablo, J. J. DNA shape dominates sequence affinity in nucleosome formation. Phys. Rev. Lett. (2014). doi:10.1103/PhysRevLett.113.168101

18. Galande, S., Purbey, P. K., Notani, D. & Kumar, P. P. The third dimension of gene regulation: organization of dynamic chromatin loopscape by SATB1. Current Opinion in Genetics and Development (2007). doi:10.1016/j.gde.2007.08.003

19. Nakagomi, K., Kohwi, Y., Dickinson, L. a & Kohwi-Shigematsu, T. A novel DNA-binding motif in the nuclear matrix attachment DNA-binding protein SATB1. Mol. Cell. Biol. (1994). doi:10.1128/MCB.14.3.1852.Updated

20. Fessing, M. Y. et al. P63 regulates Satb1 to control tissue-specific chromatin remodeling during development of the epidermis. J. Cell Biol. (2011). doi:10.1083/jcb.201101148

21. Han, H. J., Russo, J., Kohwi, Y. & Kohwi-Shigematsu, T. SATB1 reprogrammes gene expression to promote breast tumour growth and metastasis. Nature (2008). doi:10.1038/nature06781

22. Alvarez, J. D. et al. The MAR-binding protein SATB1 orchestrates temporal and spatial expression of multiple genes during T-cell development. Genes Dev. (2000). doi:10.1101/gad.14.5.521

23. Cai, S., Lee, C. C. & Kohwi-Shigematsu, T. SATB1 packages densely looped, transcriptionally active chromatin for coordinated expression of cytokine genes. Nat. Genet. (2006). doi:10.1038/ng1913

24. Balamotis, M. A. et al. Satb1 Ablation Alters Temporal Expression of Immediate Early Genes and Reduces Dendritic Spine Density during Postnatal Brain Development. Mol. Cell. Biol. (2012). doi:10.1128/MCB.05917-11

25. Agrelo, R. et al. SATB1 Defines the Developmental Context for Gene Silencing by Xist in Lymphoma and Embryonic Cells. Dev. Cell (2009). doi:10.1016/j.devcel.2009.03.006

26. Savarese, F. et al. Satb1 and Satb2 regulate embryonic stem cell differentiation and Nanog expression. Genes Dev. (2009). doi:10.1101/gad.1815709

27. Kitagawa, Y. et al. Guidance of regulatory T cell development by Satb1-dependent super-enhancer establishment. Nat. Immunol. (2017). doi:10.1038/ni.3646

28. Dickinson, L. A., Joh, T., Kohwi, Y. & Kohwi-Shigematsu, T. A tissue-specific MAR SAR DNA-binding protein with unusual binding site recognition. Cell (1992). doi:10.1016/0092-8674(92)90432-C

29. Dickinson, L. A., Dickinson, C. D. & Kohwi-Shigematsu, T. An atypical homeodomain in SATB1 promotes specific recognition of the key structural element in a matrix attachment region. J. Biol. Chem. (1997). doi:10.1074/jbc.272.17.11463

30. Bode, J. et al. Biological significance of unwinding capability of nuclear matrix-associating DNAs. Science (1992). doi:10.1126/science.1553545

31. Kohwi-Shigematsu, T. & Kohwi, Y. Torsional Stress Stabilizes Extended Base Unpairing in Suppressor Sites Flanking Immunoglobulin Heavy Chain Enhancer. Biochemistry (1990). doi:10.1021/bi00493a009

32. Purbey, P. K. et al. PDZ domain-mediated dimerization and homeodomain-directed specificity are required for high-affinity DNA binding by SATB1. Nucleic Acids Res. (2008). doi:10.1093/nar/gkm1151

33. Yamasaki, K., Akiba, T., Yamasaki, T. & Harata, K. Structural basis for recognition of the matrix attachment region of DNA by transcription factor SATB1. Nucleic Acids Res. (2007). doi:10.1093/nar/gkm504

34. Yamasaki, K. & Yamasaki, T. The combination of sequence-specific and nonspecific DNA-binding modes of transcription factor SATB1. Biochem. J. 473, 3321–3339 (2016).

35. Stasevich, T. J., Mueller, F., Brown, D. T. & McNally, J. G. Dissecting the binding mechanism of the linker histone in live cells: An integrated FRAP analysis. EMBO J. (2010). doi:10.1038/emboj.2010.24

36. Cardarelli, F. & Gratton, E. In vivo imaging of single-molecule translocation through nuclear pore complexes by pair correlation functions. PLoS One (2010). doi:10.1371/journal.pone.0010475

37. Paakinaho, V. et al. Single-molecule analysis of steroid receptor and cofactor action in living cells. Nat. Commun. (2017). doi:10.1038/ncomms15896

38. Chen, J. et al. Single-molecule dynamics of enhanceosome assembly in embryonic stem cells. Cell (2014). doi:10.1016/j.cell.2014.01.062

39. Landt, S. G. et al. ChIP-seq guidelines and practices of the ENCODE and modENCODE consortia. Genome Research (2012). doi:10.1101/gr.136184.111

40. Buenrostro, J. D., Giresi, P. G., Zaba, L. C., Chang, H. Y. & Greenleaf, W. J. Transposition of native chromatin for fast and sensitive epigenomic profiling of open chromatin, DNA-binding proteins and nucleosome position. Nat. Methods (2013). doi:10.1038/nmeth.2688

41. Tokunaga, M., Imamoto, N. & Sakata-Sogawa, K. Highly inclined thin illumination enables clear single-molecule imaging in cells. Nat. Methods (2008). doi:10.1038/nmeth1171

42. Bailey, T. L. et al. MEME Suite: Tools for motif discovery and searching. Nucleic Acids Res. (2009). doi:10.1093/nar/gkp335

43. Wang, Z. et al. The structural basis for the oligomerization of the N-terminal domain of SATB1. Nucleic Acids Res. (2012). doi:10.1093/nar/gkr1284

44. Soufi, A. et al. Pioneer transcription factors target partial DNA motifs on nucleosomes to initiate reprogramming. Cell (2015). doi:10.1016/j.cell.2015.03.017

45. Schep, A. N. et al. Structured nucleosome fingerprints enable high-resolution mapping of chromatin architecture within regulatory regions. Genome Res. (2015). doi:10.1101/gr.192294.115

46. Lowary, P. T. & Widom, J. New DNA sequence rules for high affinity binding to histone octamer and sequence-directed nucleosome positioning. J. Mol. Biol. (1998). doi:10.1006/jmbi.1997.1494

47. Zhou, T. et al. DNAshape: a method for the high-throughput prediction of DNA structural features on a genomic scale. Nucleic Acids Res. (2013). doi:10.1093/nar/gkt437

48. El Hassan, M. A. & Calladine, C. R. Propeller-twisting of base-pairs and the conformational mobility of dinucleotide steps in DNA. J. Mol. Biol. (1996). doi:10.1006/jmbi.1996.0304

49. Kimura, K. & Hirano, T. ATP-dependent positive supercoiling of DNA by 13S condensin: A biochemical implication for chromosome condensation. Cell (1997). doi:10.1016/S0092-8674(00)80524-3

50. Dror, I., Golan, T., Levy, C., Rohs, R. & Mandel-Gutfreund, Y. A widespread role of the motif environment in transcription factor binding across diverse protein families. Genome Res. (2015). doi:10.1101/gr.184671.114

51. Braeckmans, K. et al. Line FRAP with the confocal laser scanning microscope for diffusion measurements in small regions of 3-D samples. Biophys. J. (2007). doi:10.1529/biophysj.106.099838

52. Mueller, F., Wach, P. & McNally, J. G. Evidence for a common mode of transcription factor interaction with chromatin as revealed by improved quantitative fluorescence recovery after photobleaching. Biophys. J. (2008). doi:10.1529/biophysj.107.123182

53. Hajjoul, H. et al. High-throughput chromatin motion tracking in living yeast reveals the flexibility of the fiber throughout the genome. Genome Res. (2013). doi:10.1101/gr.157008.113

54. Lucas, J. S., Zhang, Y., Dudko, O. K. & Murre, C. 3D trajectories adopted by coding and regulatory DNA elements: First-passage times for genomic interactions. Cell (2014). doi:10.1016/j.cell.2014.05.036

55. Weber, S. C., Spakowitz, A. J. & Theriot, J. A. Bacterial chromosomal loci move subdiffusively through a viscoelastic cytoplasm. Phys. Rev. Lett. (2010). doi:10.1103/PhysRevLett.104.238102

56. Struhl, K. & Segal, E. Determinants of nucleosome positioning. Nature Structural and Molecular Biology (2013). doi:10.1038/nsmb.2506

57. Mirny, L. A. Nucleosome-mediated cooperativity between transcription factors. Proc. Natl. Acad. Sci. (2010). doi:10.1073/pnas.0913805107

58. Li, M. Z. & Elledge, S. J. SLIC: A method for sequence‐ and ligation-independent cloning. Methods Mol. Biol. (2012). doi:10.1007/978-1-61779-564-0_5

59. Sanjana, N. E. et al. A transcription activator-like effector toolbox for genome engineering. Nat. Protoc. (2012). doi:10.1038/nprot.2011.431

60. Ran, F. A. et al. Double nicking by RNA-guided CRISPR cas9 for enhanced genome editing specificity. Cell (2013). doi:10.1016/j.cell.2013.08.021

61. Grigoryev, S. A., Arya, G., Correll, S., Woodcock, C. L. & Schlick, T. Evidence for heteromorphic chromatin fibers from analysis of nucleosome interactions. Proc. Natl. Acad. Sci. (2009). doi:10.1073/pnas.0903280106

62. Correll, S. J., Schubert, M. H. & Grigoryev, S. A. Short nucleosome repeats impose rotational modulations on chromatin fibre folding. EMBO J. (2012). doi:10.1038/emboj.2012.80

63. Johnson, D. S., Mortazavi, A., Myers, R. M. & Wold, B. Genome-wide mapping of in vivo protein-DNA interactions. Science (80-.). (2007). doi:10.1126/science.1141319

64. Zhang, Y. et al. Model-based analysis of ChIP-Seq (MACS). Genome Biol. (2008). doi:10.1186/gb-2008-9-9-r137

65. Love, M. I., Huber, W. & Anders, S. Moderated estimation of fold change and dispersion for RNA-seq data with DESeq2. Genome Biol. (2014). doi:10.1186/s13059-014-0550-8

66. Heinz, S. et al. Simple Combinations of Lineage-Determining Transcription Factors Prime cis-Regulatory Elements Required for Macrophage and B Cell Identities. Mol. Cell (2010). doi:10.1016/j.molcel.2010.05.004

67. Denny, S. K. et al. Nfib Promotes Metastasis through a Widespread Increase in Chromatin Accessibility. Cell (2016). doi:10.1016/j.cell.2016.05.052

